# GENESIS: Generating scRNA-Seq data from Multiome Gene Expression

**DOI:** 10.1101/2025.05.06.652399

**Authors:** Simone G. Riva, Brynelle Myers, Francesca M. Buffa, Andrea Tangherloni

**Affiliations:** MRC Molecular Haematology Unit, MRC Weatherall Institute of Molecular Medicine, Oxford University, Oxford, UK; Sir William Dunn School of Pathology, Oxford University, Oxford, UK; Department of Computing Sciences, Bocconi Institute for Data Science and Analytics, Bocconi University, Milan, Italy

**Keywords:** Generative AI, GANs, VAEs, scRNA-Seq, Multiome Gene Expression

## Abstract

Single-cell technologies have significantly advanced our understanding of cellular heterogeneity by allowing the examination of individual cells at high resolution. Traditional single-cell RNA sequencing (scRNA-Seq) methods, which utilise whole cells, capture comprehensive RNA content. In contrast, emerging Multiome technologies, which simultaneously profile multiple omics such as gene expression (GEX) and chromatin accessibility, rely on nuclear RNA, potentially missing key cytoplasmic information. This discrepancy leads to substantial technical and biological differences between GEX and scRNA-Seq datasets, making it difficult to integrate data and perform downstream tasks like cell-type classification. To address this challenge, we introduce GENESIS (Gene Expression Normalisation and Enhancement for Single-cell Integrated Sequencing), a novel computational framework designed to transform GEX data from Multiome experiments into enhanced, scRNA-Seq like profiles. Utilising advanced generative models—including Variational Autoencoders, Generative Adversarial Networks, and a tailored VAE UNet architecture—GENESIS can generate high-quality data by modelling and compensating for the inherent differences between nuclear and cytoplasmic RNA. Our comprehensive evaluations show that GENESIS, particularly through the VAE UNet model, generates synthetic scRNA-Seq data that closely resembles the resolution and biological accuracy of whole-cell sequencing, improving downstream tasks, especially cell-type classification.

## I. Introduction & motivation

Single-cell technologies have revolutionised our understanding of cellular heterogeneity and biological systems by enabling researchers to examine individual cells at unprecedented resolution [1], [2]. Unlike traditional bulk sequencing methods that average signals across thousands of cells, single-cell approaches provide detailed insights into cellular diversity, rare cell populations, and dynamic biological processes at the individual cell level [3]. Over the past decade, significant technological advances have led to the development of various single-cell analysis platforms, with single-cell RNA sequencing (scRNA-Seq) emerging as a particularly powerful tool [4], [5]. These technologies have transformed multiple fields, including developmental biology, immunology, cancer research, and neuroscience, by revealing previously undetectable cell-to-cell variations, as well as helping to identify new cell types and states [6], [7].

The ability to profile individual cells has unveiled complex cellular hierarchies, developmental trajectories, and regulatory networks that were previously obscured in bulk analyses [8], [9]. This granular view has proven especially valuable in understanding tissue heterogeneity, disease progression, and cellular response to various stimuli. In addition, single-cell technologies have enabled researchers to track cellular differentiation pathways, study rare cell populations, and investigate cellular response mechanisms at unprecedented resolution [10], [11]. However, single-cell data analysis presents unique challenges, including technical noise, dropout events, and batch effects, which require sophisticated computational methods for proper analyses and interpretation [12], [13]. The development of robust analytical tools and standardised protocols has become crucial for extracting meaningful biological insights from these complex datasets [14]–[17].

Besides that, recent advances in single-cell technologies have enabled simultaneous profiling of multiple molecular modalities within the same cell, known as Multiome or multiomics approaches [18], [19], posing further analysis challenges. These methods, such as SHARE-Seq, 10x Genomics Multiome, and CITE-Seq, allow researchers to simultaneously measure gene expression (GEX) alongside other molecular features, including chromatin accessibility, protein levels, or DNA methylation status [20], [21]. The integration of multiple data types provides a more comprehensive understanding of cellular state and regulatory mechanisms, revealing how different molecular layers interact to determine cell identity and function [22]. For instance, paired measurements of chromatin accessibility and gene expression can illuminate the relation-ship between regulatory element activity and transcriptional output, providing insights into gene regulation mechanisms that would be impossible to obtain from either measurement alone [23]. However, the analysis of multimodal data presents additional computational challenges, requiring sophisticated methods for data integration, normalisation, and interpretation [24]. Moreover, a fundamental technical distinction between traditional scRNA-Seq and Multiome approaches lies in their starting biological material. Standard scRNA-Seq typically uses whole cells, allowing access to both nuclear and cytoplasmic RNA content, thus capturing a more complete snapshot of cellular RNA content [25]. In contrast, Multiome technologies, particularly those involving chromatin accessibility measurements (GEX and scATAC-Seq), require intact nuclei rather than whole cells [26]. This requirement for nucleus isolation means that cytoplasmic RNA is largely lost during sample preparation, potentially reducing the RNA detection sensitivity compared to whole-cell approaches [27]. The nuclear isolation procedure also impacts the RNA species that can be detected, as only nuclear-localised and nascent transcripts are readily accessible, while mature cytoplasmic mRNAs are largely excluded [28]. This distinction is crucial for experimental design and data interpretation, as nuclear RNA represents only about 20-50% of the total cellular RNA content and may provide a different view of the transcriptional landscape of the isolated cells compared to whole-cell measurements [29], [30].

Despite significant advances in single-cell transcriptomics, genomics, and computational methods, neither computational nor machine learning tools currently exist to enhance nuclear RNA data from Multiome experiments to match the quality and resolution of scRNA-Seq data. While various computational methods have been developed for batch correction [31], [32] and data integration [33], [34], these approaches primarily focus on harmonising similar data types or addressing technical variations within the same modality. The fundamental differences between nuclear and cytoplasmic RNA content present unique challenges that existing tools do not explicitly address. Current methods, such as Seurat v4, LIGER, and Harmony, can integrate different modalities but do not attempt to enhance the signal quality of nuclear RNA *synthetically* to match whole-cell transcriptomes. This gap in the field represents a significant opportunity for methodological advancement, particularly as Multiome experiments become increasingly prevalent in single-cell research.

Just focusing on cell type annotation, classifiers trained on scRNA-Seq data, relying on predefined and well-known marker genes, often fail to predict equivalent cell populations GEX multiome datasets accurately. This discrepancy arises due to technical and biological differences between platforms, including variations in sensitivity, gene detection rates, and integration with other modalities, such as chromatin accessibility. As a result, models that perform well on scRNA-Seq data may not generalise to multiome data, leading to misclassification or an inability to detect certain cell populations reliably. In this study, we thus present GENESIS (Gene Expression Normalisation and Enhancement for Single-cell Integrated Sequencing), a comprehensive computational framework designed to address the unique challenge of transforming the direct gene expression level of Multiome in scRNA-Seq data. GENESIS specifically tackles the technical variations and biological differences between whole-cell RNA-Seq and nuclear RNA-Seq data, providing a robust method for cross-platform normalisation and enhancing gene expression levels. Our approach leverages Machine Learning (ML) algorithms, accounting for the systematic differences in RNA capture efficiency between cellular and nuclear preparations, while preserving biological signals critical for downstream analyses. GENESIS uses ML techniques to model and correct for technical artefacts, enabling more accurate integration of datasets generated using different protocols. We show the utility of GENESIS through extensive validation using scRNA-Seq data from Kumar *et al*. [35], as well as Multiome data combining GEX with scATAC-Seq from Bhat-Nakshatri *et al*. [36], from matching tissue and cell types.

## II. Data

In this study, we analysed single-cell RNA sequencing and Multiome data from two key datasets: Kumar *et al*. dataset comprises single-cell RNA sequencing data from whole cells, providing comprehensive transcriptional profiles across diverse cell populations; in contrast, the Bhat-Nakshatri *et al*. dataset consists of Multiome measurements, including nuclear RNA expression and chromatin accessibility data. These datasets were obtained from the CellXGene data portal [37], a centralised repository for single-cell genomics data. Specifically, we downloaded the h5ad (Hierarchical Data Format 5 with Annotated Data) files for both datasets, which are publicly available and maintain the original data structure and annotations. The h5ad format, implemented through the AnnData Python package [38], preserves both the raw expression matrices and associated metadata, including cell annotations, quality metrics, and experimental conditions. These files serve as the foundation for our analysis pipeline, ensuring reproducibility and maintaining data provenance. The datasets can be accessed through the following CellXGene collection URLs: https://cellxgene.cziscience.com/collections/4195ab4c-20bd-4cd3-8b3d-65601277e731, https://cellxgene.cziscience.com/collections/77446b76-1c2d-4a71-8e59-0efd4374d98e, allowing other researchers to reproduce our analyses using identical input data.

## III. Method

In this work, we explored the use of generative models, especially Variational Autoencoders (VAEs) [39] and Generative Adversarial Networks (GANs) [40], to learn transformations of GEX profiles in scRNA-Seq profiles. These models are employed to map the same cell populations from GEX to scRNA-Seq data by learning structured representations of the underlying gene expression distributions. VAEs offer significant advantages over both standard autoencoders (AEs) and deterministic feedforward models, such as multilayer perceptrons (MLPs), particularly in tasks involving reconstruction, domain translation, or dataset alignment. Unlike AEs, which learn a deterministic mapping from input to latent representation without constraints, VAEs impose a probabilistic prior, typically a standard Gaussian, on the latent variables, enforcing a structured latent space by approximating the posterior distribution over latent variables and aligning it with the chosen prior. This regularisation yields a smooth, continuous, and structured latent space, which improves generalisation and enables meaningful interpolation and sampling. In contrast to AEs and MLPs that directly learn point-to-point mappings, VAEs capture the underlying data distribution and model uncertainty explicitly. This allows them to handle ambiguous or multimodal relationships between input and output domains more effectively, avoiding overfitting and promoting more robust, coherent transformations across diverse datasets. In addition to VAEs, we investigated GANs as a complementary approach due to their capacity to generate highly realistic samples by directly modelling the target data distribution. While VAEs focus on latent space regularisation and uncertainty modelling, GANs excel at refining the sharpness and realism of generated outputs by learning through adversarial feedback. Given the structured and high-dimensional nature of scRNA-Seq data, GANs could enhance reconstruction quality, particularly for genes not present in the input GEX matrix, by better capturing complex gene co-expression patterns.

We used gene expression profiles (GEX) as input to our generative models, where each input vector (i.e., a cell profile) consists of *G* genes representing a subset of the full transcriptome. The goal of our models is to reconstruct the full scRNA-Seq expression profile comprising *R* genes, where *G < R*. During training, the models learned to reconstruct the expression of the *G* scRNA-Seq genes while simultaneously generating plausible expression values for the remaining *R* − *G* genes not present in the input (i.e., GEX profile). To ensure biological consistency and facilitate supervised learning, we selected *N* cells in both sequencing experiments. In addition, during training, the input GEX and target scRNA-Seq profiles were matched by cell type. To be more precise, training was performed over mini-batches of matched GEX and scRNA-Seq pairs from the same cell types, allowing the models to learn meaningful cross-modal representations while reconstructing the observed genes and predicting the unobserved ones.

### A. GANs

To explore the generative potential of deep adversarial learning, we implemented a GAN to synthesise scRNA-Seq profiles from the lower-dimensional GEX input. The generator network consists of an initial linear layer (from *G* genes to 512 dimensions) followed by a ReLU activation, then two hidden layers with linear layer, batch normalisation, and ReLU activation, with 1024 and 2048 neurons, respectively. The final layer is a fully connected linear layer of *R* neurons, followed by a ReLU activation. The discriminator has the same architecture without batch normalisation and replaces ReLU with LeakyReLU activations (negative slope = 0.2) to maintain gradient flow during training. In this case, the final layer is just a single neuron for binary classification. Both networks were trained using the RMSprop optimiser (learning rate = 0.0001), a choice informed by prior stability analyses in GAN literature.

GANs use these two MLPs that compete against each other in a minimax game: the generator attempts to produce outputs that resemble real data, while the discriminator learns to distinguish real scRNA-Seq profiles from the generated ones. Ideally, this adversarial process leads to a generator capable of producing highly realistic synthetic data. However, when training a standard GAN in this context, we observed rapid mode collapse. After just a few epochs, the discriminator was able to easily identify fake samples and stop providing meaningful gradient updates to the generator, leading the generator to fail entirely. This outcome is expected in high-dimensional, structured biological data, where standard GAN loss functions often suffer from vanishing gradients. To address this instability, we used the Wasserstein GAN (WGAN) framework [41], which replaces the binary cross-entropy loss with a continuous approximation of the Earth Mover (Wasserstein-1) distance between real and generated distributions. This modification significantly improves training stability, as the discriminator (called the “critic” in WGAN) provides informative gradients even when its classification is near perfect. As a result, the generator received significant feedback throughout training, allowing it to approximate the target distribution better and generate realistic high-dimensional scRNA-Seq profiles from compressed GEX input. Both the generator and critic were trained using the RMSprop optimiser (learning rate = 0.00005). We clipped the critic’s weights in the range [−0.01, 0.01] to enforce the Lipschitz constraint required by the WGAN training procedure. For each batch in the training set, we trained the critic 5 times before updating the generator.

### B. VAE

The VAE architecture consists of an encoder that maps the input gene expression profile to a latent space defined by the mean and log-variance of a Gaussian distribution. A latent vector is then sampled using the reparameterisation trick and passed to the decoder, which reconstructs the full scRNA-Seq expression profile. The model is trained to minimise a loss (i.e., the ELBO) that combines a reconstruction error and a Kullback-Leibler (KL) divergence term that regularises the latent space toward a standard normal distribution [39]. In our tests, the encoder consists of an initial linear layer (from *G* genes to 2048 dimensions) followed by a ReLU activation, then two hidden layers with a linear layer, batch normalisation, and ReLU activation, with the following dimensions 1024 and 512, respectively. The last hidden layer is then mapped to two latent vectors of 32 dimensions representing the mean and log-variance of a Gaussian distribution. The decoder mirrors the encoder structure, projecting the latent vector back to the scRNA-Seq gene space using symmetric layers without batch normalisation. The final reconstruction is generated using a fully connected output layer of *R* neurons, followed by a ReLU activation. During training, the model uses the standard reparameterisation trick to sample from the latent space and optimise the ELBO function, combining Mean Squared Error (MSE) loss and a weighted KL divergence term (we set *β* = 0.1). The entire model was trained using the Adam optimiser with a learning rate of 0.0001.

### C. VAE UNet

To further enhance the ability of our generative framework, we designed an *ad hoc β*VAE UNet model tailored for high-dimensional gene expression reconstruction. This architecture builds upon a traditional VAE by integrating UNet-style skip connections between the encoder and decoder [42], allowing for a more efficient transfer of intermediate representations (see Fig. 1). The encoder consists of an initial linear layer (from *G* genes to 2048 dimensions) followed by a ReLU activation, then two residual blocks with batch normalisation and ReLU activations, with 1024 and 512 neurons, respectively. These residual blocks facilitate stable training and preserve important features at each resolution. During encoding, intermediate outputs are stored as skip connections, which are later reused by the decoder to guide the reconstruction process. The decoder mirrors the encoder’s structure without batch normalisation and residual connections and progressively integrates the skip features via concatenation at each decoding stage. This structure enables the model to recover fine-grained patterns from the input GEX data that may be lost through deep compression alone. By aligning encoder and decoder layers in this hierarchical manner, the model benefits from both global latent representations and localised gene co-expression patterns, ultimately improving the generated scRNA-Seq profiles. As for the standard *β*VAE, the final layer is a fully connected layer of *R* neurons, followed by a ReLU activation. Also in this case, we trained the model using the standard reparameterisation trick to sample from the latent space and optimise the ELBO function, combining MSE loss and a weighted KL divergence term (we set *β* = 0.1). The entire model was trained using the Adam optimiser with a learning rate of 0.0001.

**Figure 1.**
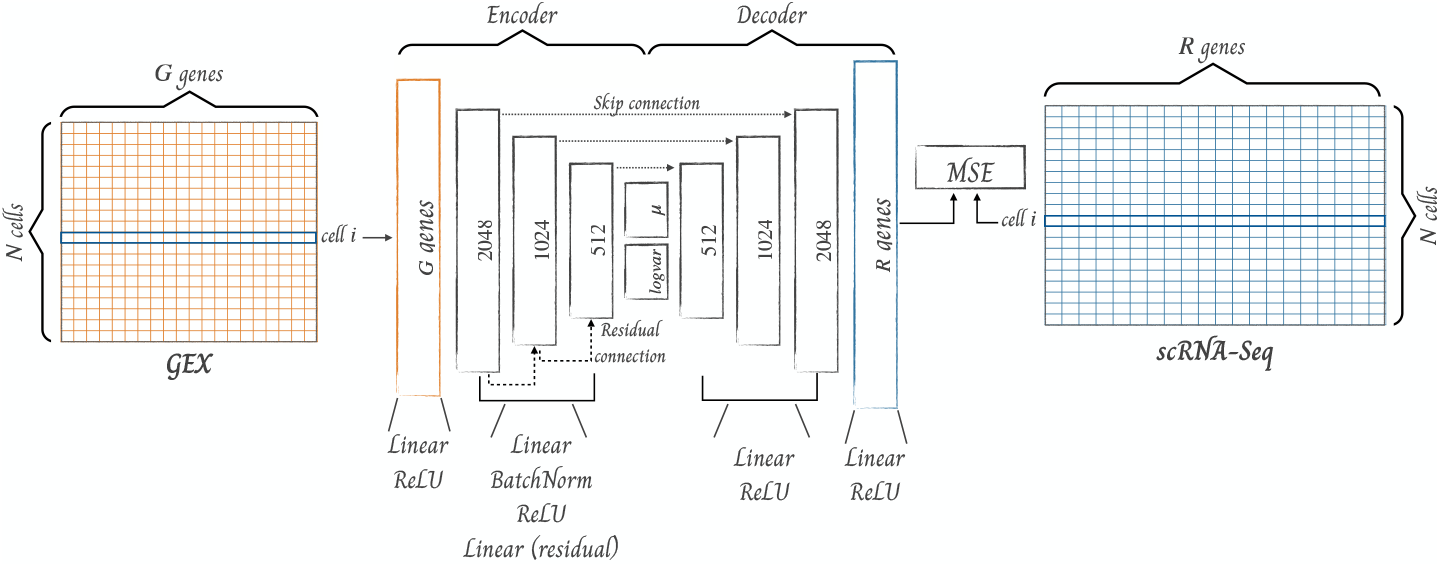
The VAE UNet architecture is composed of an encoder that projects the input gene expression profile (i.e., cell *i* of the GEX) into a latent space parameterised by a mean and log-variance, followed by a decoder that reconstructs a cell, matching the cell *i* of the scRNA-Seq profile, from a sampled latent vector, exploiting skip connections from the encoder’s layers. Training minimises a combined loss of MSE and KL divergence, encouraging the latent space to follow a smooth, structured distribution for better generalisation.

We designed this VAE UNet architecture to address the limitations of standard VAEs in reconstructing sparse, structured expression data. In particular, skip connections allow the decoder to reconstruct the observed GEX genes better while improving the generation of the unobserved gene subset, especially when input-output mappings are complex and information-rich. This design choice reflects the biological intuition that gene regulation is hierarchical and context-dependent and enables the model to synthesise expression profiles that are both accurate and biologically coherent.

## IV. Result

Since the Bhat-Nakshatri *et al*. dataset contains 51, 367 single cells, whereas the Kumar *et al*. dataset contains 714, 331 single cells (∼ 14×), firstly, we carefully renamed cell type identities to map their names in both datasets correctly — considering the following five cell types: basal cell, fibroblast, endothelial cell, T cell, and macrophage, since identified in both datasets — and secondly, performed a strategic subsetting on both dataset. The first subsetting, applied on Kumar *et al*. dataset, was implemented based on specific cell annotations (listed above) and the stored metadata, specifically on the used assay, selecting only the 10x 3’ v3 sequencing assay method, leaving us with 385, 614 cells. To ensure balanced representation across cell types, we performed a second round of subsetting based on cell identity distributions. We first determined the minimum number of cells per identity type present across both datasets. Then, we randomly sampled cells from each identity category to match these minimum counts, resulting in balanced datasets of 31, 650 cells each. The final composition of each dataset comprised 16, 184 basal cells, 6, 526 fibroblasts, 5, 407 endothelial cells, 2, 240 T cells, and 1, 293 macrophages. This balanced subsetting approach eliminated potential biases that could arise from uneven cell type distributions between the datasets. To confirm that after the applied subsetting strategies and cell types mapping, we utilised six marker genes for each cell type (within our mapped cell types) previously identified in Kumar *et al*., which served as a reference point to confirm cell identity in our analysis. We generated a matrix plot to visualise the expression patterns of these marker genes across the mapped cell populations in our subsetted scRNA-Seq data (see Fig. 2). The visualisation revealed distinct gene expression signatures that uniquely identified each cell population, with strong expression of cell type-specific markers and minimal cross-population expression, as shown in Fig. 1d in [35]. This analysis confirmed that these canonical markers showed high expression levels in their expected cell types: *IL7R, CCL5, PTPRC, CXCR4, GNLY, CD2*, and *SRGN* were enriched in T cells, *KRT14, KRT17, DST, KRT5, SAA1, ACTA2*, and *SFN* in basal cells, *SELE, ACKR1, FABP4, STC1, ANGPT2*, and *CSF3* in endothelial cells, *DCN, APOD, CFD, TNFAIP6, LUM, COL1A2*, and *COL1A1* in fibroblasts, and *HLA-DRA, IL1B, HLA-DPA1, HLA-DPB1, HLA-DRB1, CD74*, and *CCL3* in macrophages. This expression pattern closely mirrored the findings reported in Kumar *et al*., validating and confirming our cell type mapping. In contrast, when we applied the same marker gene analysis to the subsetted Multiome GEX dataset, we observed notably different expression patterns. While these established markers effectively delineated cell populations in the scRNA-Seq data, their expression patterns in the nuclear RNA data were less distinct. This difference can be attributed to the inherent characteristics of nuclear RNA sequencing [28]. The matrix plot of the GEX dataset (Fig. 3) revealed more diffuse expression patterns of the marker genes, with reduced signal intensity and increased noise compared to the scRNA-Seq data.

**Figure 2.**
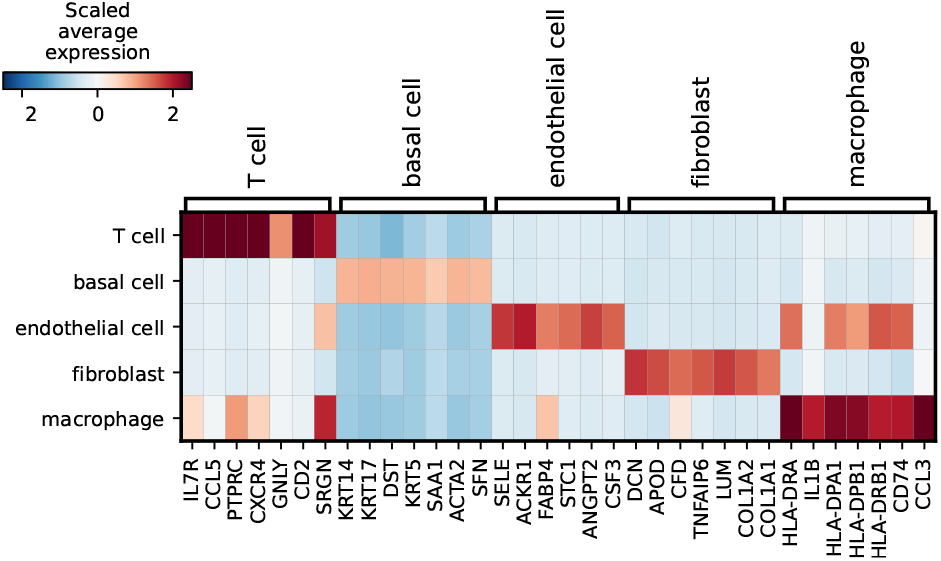
Matrix plot showing scaled expression of cell type-specific marker genes across mapped cell populations in the subsetted scRNA-Seq data. The heatmap displays canonical markers for T cells, basal cells, endothelial cells, fibroblasts, and macrophages. Red colours indicate high expression, while blue colours represent low expression. The plot demonstrates distinct expression patterns of these markers across their corresponding cell types, validating cell type annotations. The colour scale represents scaled average expression from -2 (blue) to 2 (red).

**Figure 3.**
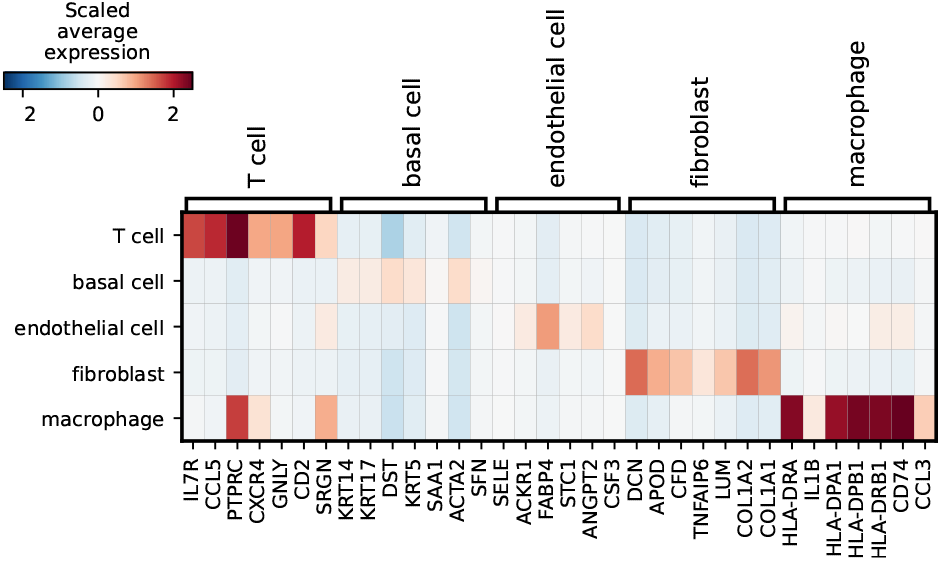
Matrix plot showing scaled expression of canonical marker genes in nuclear RNA (GEX) data across mapped cell populations. The heatmap reveals less distinct expression patterns compared to scRNA-Seq, with T cell, basal cell, endothelial cell, fibroblast, and macrophage markers. The more diffuse expression patterns and reduced signal intensity are characteristic of nuclear RNA sequencing. The colour scale represents scaled average expression from -2 (blue) to 2 (red), highlighting the challenges in cell type identification using established markers in nuclear RNA data.

To quantitatively assess the differences in marker gene expression between the subsetted scRNA-Seq and Multiome GEX datasets, we generated violin plots comparing the expression distributions across cell types in Fig. 4. These plots provided a direct visualisation of expression level differences between the two modalities for each marker gene within their corresponding mapped cell populations. The violin plots revealed consistently higher expression levels and better signal-to-noise ratios in the scRNA-Seq data compared to the GEX data.

**Figure 4.**
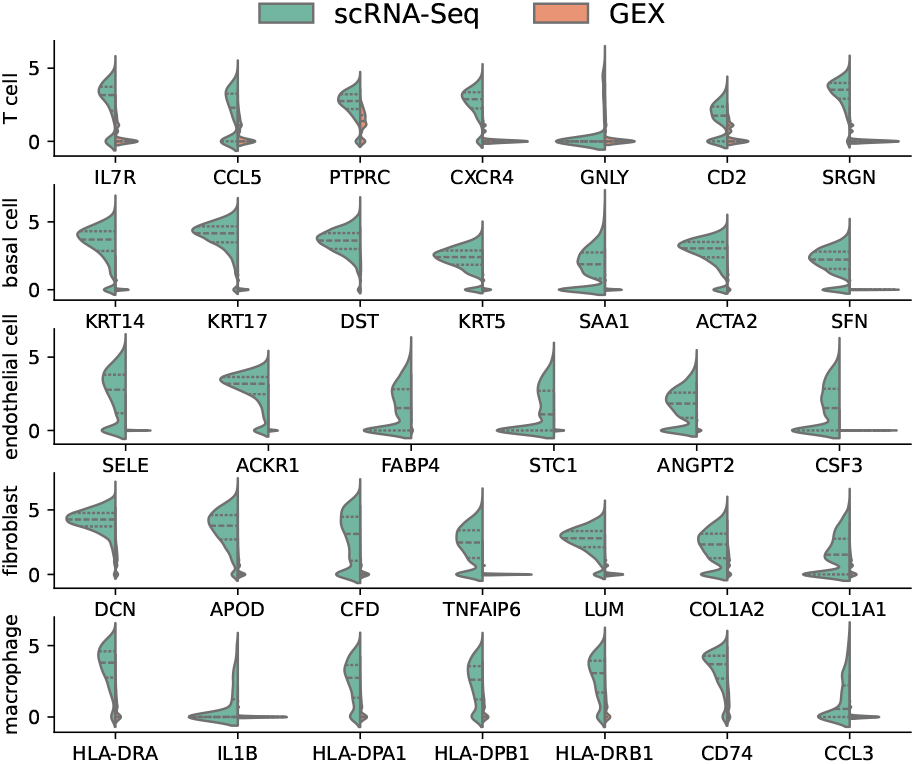
Violin plots comparing gene expression distributions between scRNA-Seq (green) and GEX (orange) data across different cell types. The plots display cell type-specific markers for T cells, basal cells, endothelial cells, fibroblasts, and macrophages. The comparison demonstrates systematic differences in gene detection and expression levels between whole-cell and nuclear RNA sequencing approaches, with scRNA-Seq generally showing higher sensitivity and more defined expression patterns. The Y-axis represents log-normalised expression values, and the width of each violin indicates the density of cells at each expression level.

To set a robust and interpretable baseline classifier for cell type classification, we trained an XGBoost model on scRNA-Seq data using the previously reported curated set of well-established marker genes, resulting in a compact and biologically meaningful feature set. This targeted approach allowed us to reduce noise and complexity while retaining the key discriminative signals necessary for accurate classification. We chose XGBoost [43] for its strong performance with structured data and its inherent interpretability, which is especially valuable when working with a small number of biologically validated features. The model provides feature importance metrics that can help validate or refine existing marker gene selections, offering both predictive power and biological insight. Hyperparameter optimisation was performed using a grid search with 5-fold cross-validation to ensure the generalizability and stability of the model. The parameter grid

included the following hyperparameters.

- booster type: `gbtree` and `dart`;
- maximum tree depth: 2, 4, and 6;
- learning rate: 0.01, 0.05, and 0.1;
- number of estimators 50, 100, and 150;
- subsampling rate 0.6, 0.8, and 1.0.

The optimal configuration identified through this process was: `dart` booster, learning rate equal to 0.1, max depth equal to 4, 150 estimators, and subsampling rate equal to 0.6.

To evaluate and compare the performance of XGBoost on the five datasets (real scRNA-Seq, real GEX, and generated GEX with VAE, GAN, and VAE UNet), we used the F1 Score (eq. 1) as our primary metric.

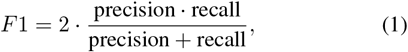

where precision (eq. 2) and recall (eq. 3) are defined as:

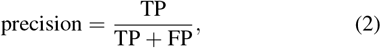

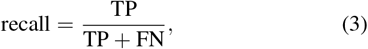

where TP = True Positives, FP = False Positives, and FN = False Negatives. The F1 Score was chosen as it provides a balanced measure of model performance by combining both precision and recall into a single metric, making it particularly suitable for our mapped cell-type specific task. This metric is especially valuable in our context as it accounts for both false positives and false negatives, considering the unbalanced number of cells per cell type. The F1 Score ranges from 0 to 1, where 1 indicates perfect precision and recall, allowing us to effectively quantify how well each model preserves cell type-specific features in the generated data. This metric is particularly relevant for single-cell analysis as it helps ensure that the synthetic data maintains both the specificity of cell type markers (precision) and the completeness of cell type-specific expression patterns (recall), which are essential for downstream biological interpretation and analysis.

Based on our comprehensive evaluation metrics — shown in Fig. 5 — the VAE UNet architecture demonstrates superior performance in generating synthetic single-cell data compared to both VAE and WGAN approaches. The VAE UNet significantly outperforms other architectures, achieving an F1 score of 0.9717. This performance nearly matches the quality of real scRNA-Seq data, representing a substantial improvement over the WGAN (F1=0.7938) and traditional VAE (F1=0.3460) architectures. The excellent performance of VAE UNet can be attributed to its hybrid architecture, which combines the generative capabilities of VAEs with the precise feature extraction of UNet structures. These results suggest that VAE UNet provides a more robust and accurate framework for generating synthetic single-cell data, potentially offering new opportunities for data augmentation and analysis in single-cell research.

**Figure 5.**
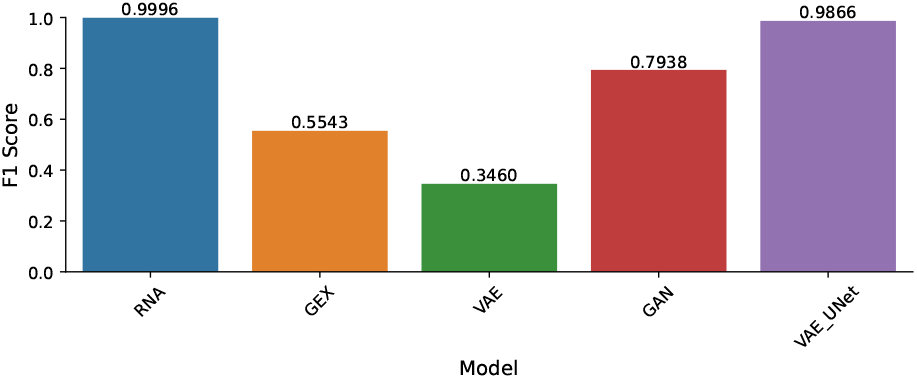
Comparison of F1 scores using scRNA-Seq, GEX, and generated data by GAN, VAE, and VAE UNet. Using scRNA-Seq data, the trained XGBoost achieved the highest F1 score (0.9996), representing the ground truth performance. XGBoost showed poor performance using GEX (F1=0.5543), reflecting the inherent challenges of nuclear RNA sequencing. Among the generative models, VAE UNet demonstrates superior performance. Indeed, XGBoost obtained a high F1 score (F1=0.9866), which is close to the F1 score achieved using scRNA-Seq data. Data generated using WGAN allowed for reaching an F1 score equal to 0.7938, while the basic VAE generated poor-quality data (F1=0.3460). This comparison clearly illustrates the effectiveness of our VAE UNet model in generating high-quality synthetic single-cell data that closely resembles the characteristics of whole-cell RNA sequencing data.

Having identified VAE UNet as the best model for synthetic data reconstruction, we evaluated its performance through detailed marker gene analysis. Fig. 6 reports the expression patterns of canonical cell-type markers using the data generated by the trained VAE UNet. This matrix plot shows that these synthetic data maintain distinct cell-type-specific expression signatures, closely mirroring the patterns observed in the original scRNA-Seq data (Fig. 2).

**Figure 6.**
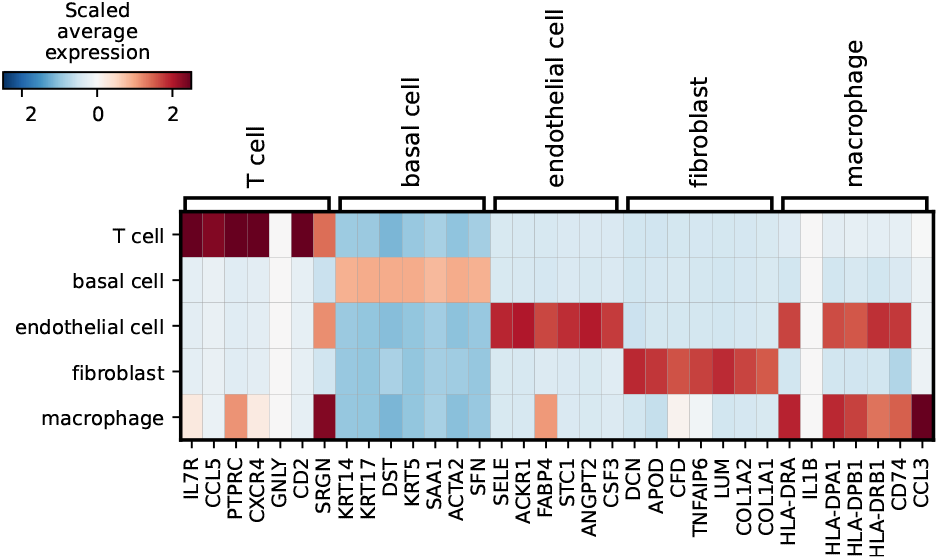
Matrix plot showing scaled expression of cell type-specific marker genes in VAE UNet generated data across the cell populations. The heatmap shows that the synthetic data maintains distinct cell type-specific expression patterns within T cells, basal cells, endothelial cells, fibroblasts, and macrophage markers. Red colours indicate high expression while blue colours represent low expression (scale from -2 to 2). The clear separation of marker gene expression patterns across different cell types suggests that the VAE UNet successfully preserves cell type-specific transcriptional signatures in the generated data, closely resembling the patterns observed in the original scRNA-Seq data shown in Fig. 2.

To further validate the quality of the generated data, we created violin plots comparing the expression distributions of key marker genes between the original scRNA-Seq and VAE UNet generated data (see Fig. 7). These plots indicate highly similar expression patterns and distributions across all examined cell types, with the synthetic data accurately capturing both the magnitude and variability of gene expression observed in the original scRNA-Seq data. The key markers used for T cells, basal cells, endothelial cells, fibroblasts, and macrophages in Kumar *et al*. paper showed remarkably consistent expression patterns between the original and synthetic datasets, further confirming the quality of our VAE UNet model.

**Figure 7.**
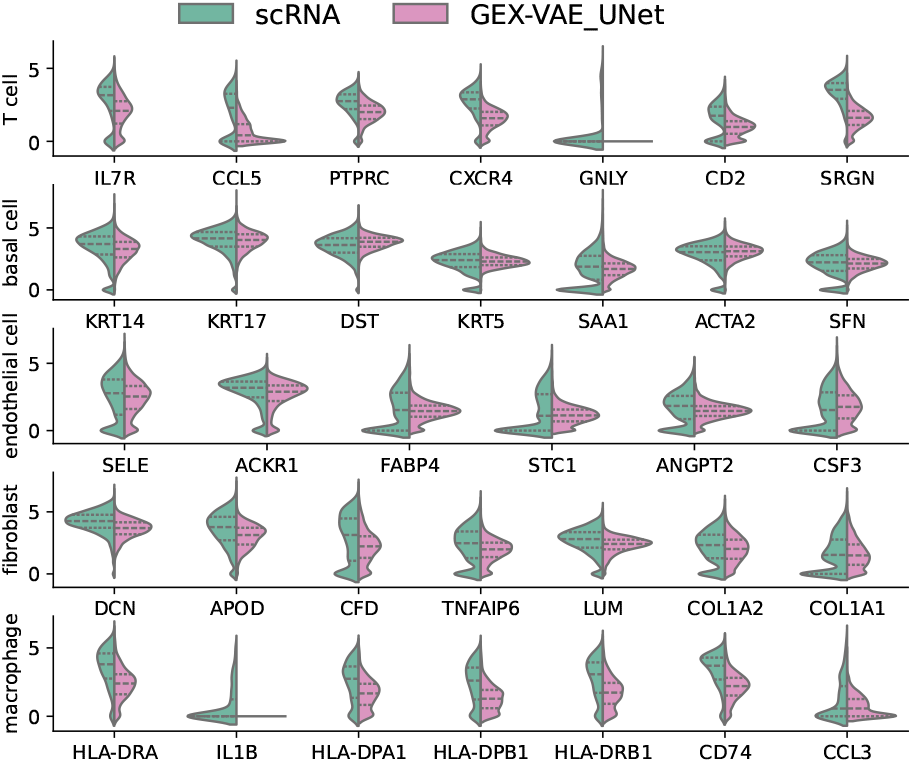
Violin plots comparing gene expression distributions between original scRNA-Seq data (green) and VAE UNet generated data (pink) across different cell types and marker genes. The plots show highly concordant expression patterns for cell type-specific markers. The similar shapes and distributions of the violin plots demonstrate that the VAE UNet generated data successfully captures both the magnitude and variability of gene expression observed in the original scRNA-Seq data, showing the model’s ability to generate biologically meaningful synthetic data.

## V. Conclusion & future work

In this study, we introduced GENESIS, a novel computational framework designed to address the challenges of integrating and analysing single-cell Multiome data. Through a comprehensive evaluation of different deep learning architectures, we showed that our VAE UNet model significantly outperforms traditional approaches in generating synthetic single-cell data. To evaluate the performance of the generative models, we trained a XGBoost classifier using on scRNA-Seq to predict the cell populations. XGBoost achieved an F1 score of 0.9866 using the data generated with VAE UNet, nearly matching the performance observed considering the scRNA-Seq data used for training the model (i.e., F1 score equal to 0.9996), while it obtained a F1 score equal to 0.7938 and 0.3460 using the data generated by WGAN and VAE, respectively. We further evaluated the quality of the data generated by VAE UNet through a specified analysis of cell type-specific marker genes, showing that VAE UNet can reconstruct the expression patterns and distributions across diverse cell populations.

Our results have important implications for single-cell transcriptomics and genomics, particularly in Multiome data analysis. The ability of GENESIS to generate high-quality synthetic data that closely mirrors the characteristics of original scRNA-Seq data while accounting for the unique properties of nuclear RNA provides new opportunities for data augmentation and integration in single-cell studies. This capability is especially valuable given the increasing prevalence of Multiome experiments and the persistent technical challenges in comparing data across different molecular modalities. Future applications of GENESIS could facilitate a more robust integration of single-cell datasets, enable a better understanding of cellular heterogeneity, and support the development of more comprehensive cell atlases. Moreover, the framework’s ability to generate synthetic data could help address issues of data sparsity and noise that often complicate single-cell analyses.

Several promising directions exist for extending the capabilities of GENESIS and broadening its applications. For example, it could be extended to analyse other single-cell modalities like scATAC-Seq.

## Acknowledgment

S.G.R. is supported by the MRC grant (MC UU 00029/3).

